# Genome sequence and characterization of a hypervirulent BI/NAP1/027 *Clostridioides difficile* (CDC20121308)

**DOI:** 10.1101/2024.04.25.590960

**Authors:** Laureano A. Español, Miranda C. Palumbo, Angela M. Barbero, Sabina Palma, Federico Serral, Diego Ruggeri, Marcelo A. Martí, Darío Fernández Do Porto, Rodrigo E. Hernández Del Pino, Virginia Pasquinelli

**Affiliations:** Centro de Investigaciones Básicas y Aplicadas (CIBA), Universidad Nacional del Noroeste de la Provincia de Buenos Aires (UNNOBA). B6000DNE, Buenos Aires, Argentina; Instituto de Cálculo, Facultad de Ciencias Exactas y Naturales, Universidad de Buenos Aires. C1428EGA, Buenos Aires, Argentina; Departamento de Química Biológica, Facultad de Ciencias Exactas y Naturales, Universidad de Buenos Aires. C1428EGA, Buenos Aires, Argentina; Centro de Investigaciones y Transferencias del Noroeste de la Provincia de Buenos Aires (CIT NOBA), UNNOBA-Universidad Nacional de San Antonio de Areco (UNSAdA)-Consejo Nacional de Investigaciones Científicas y Técnicas (CONICET). B2700, Buenos Aires, Argentina; Departamento de Bacteriología, Instituto Nacional de Enfermedades Infecciosas (INEI). ANLIS “Dr. Carlos G. Malbrán”. C1282AFF, Buenos Aires, Argentina; Instituto de Química Biológica de la Facultad de Ciencias Exactas y Naturales (IQUIBICEN) CONICET, Pabellón 2 de Ciudad Universitaria, Ciudad de Buenos Aires C1428EGA, Argentina

**Author notes:** These authors share first authorship (Laureano A. Español, Miranda C. Palumbo). These authors contributed equally to this work and share second authorship (Angela M. Barbero, Sabina Palma). These authors contributed equally to this work and share last authorship. ^#^ (Rodrigo E. Hernández Del Pino, Virginia Pasquinelli). **Correspondence:** Pasquinelli, Virginia, Hernández Del Pino, Rodrigo E.

**Keywords:** *Clostridioides difficile*_1_, NAP1/BI/027_2_, Whole Genome Sequencing_3_, Antibiotic Resistance_4_, Hypervirulent Strain_5_

## Abstract

*Clostridioides difficile* is a gram-positive bacterium implicated in antibiotic-associated diarrhea. The use of antibiotics alters the gut microbiota, rendering the host susceptible to infection by *C. difficile.* This pathogen colonizes the large intestine of humans and animals leading to asymptomatic carriage or clinical manifestations such as toxic megacolon and fulminant colitis depending on a wide range of pathogen and host factors. The emergence of BI/NAP1/027 strains in North America and the spread of these hypervirulent ribotypes worldwide have been linked to the increase in incidence and severity of *C. difficile* infections (CDI) over the last decade. In this work, we aimed to characterize the BI/NAP1/027 *C. difficile* commercial strain CDC20121308 widely employed in the study of host-pathogen interactions.

The genome sequence was obtained using a whole-genome shotgun strategy. A total of 3,717 coding sequences (CDS) and 45 tRNAs were predicted. The annotation of the CDC20121308 strain identified 26% of CDS into RAST subsystems. We also detected the presence of RT 027 lineage markers such as *thyA, cdtA, cdtB and tc*dC 18bp-deletion. Moreover, the genome of CDC20121308 had 11 genes devoted to resistance to toxic compounds, antibiotics (e.g. Tetracycline (Tet) and Vancomycin (Van)) and disinfecting agents as predicted using CARD. *C. difficile* CDC20121308 resistance to Van and Tet was confirmed by broth microdilution assay. Crystal violet staining demonstrated biofilm formation, which could be associated with antibiotic resistance and pathogenicity. Additionally, we observed a spreading diffuse growth in soft agar tubes, suggesting a motile phenotype. Lastly, a genomic region containing Type 4 Secretory System components such as virD4, virB4, and virB6 was identified.

In conclusion, our results allowed a genomic and functional characterization of the BI/NAP1/027 *C. difficile* CDC20121308 strain. We demonstrated the presence of several genes associated with pathogenesis that were validated by experimental assays. This study provides additional data for the use of this highly virulence commercial strain of epidemiological relevance in research works involving *in vitro* and *in vivo* approaches.

## Introduction

*Clostridioides difficile* (*C. difficile,* formerly *Clostridium difficile*) is a gram-positive, obligate anaerobic and spore forming bacterium widespread in the environment. *C. difficile* infection (CDI) causes a spectrum of disease, ranging from occasional diarrhea to colitis, toxic megacolon, and potentially death; which could be mistaken with other medical conditions. CDI is the major cause of antibiotic-associated diarrhea and pseudomembranous colitis in United States (Curcio et al., 2019a). Indeed, CDI ranks above methicillin resistant *Staphylococcus aureus* (MRSA) as the leading cause of hospital-acquired infection (Meyer et al., 2012; Peterson et al., 2020; Public Health England (PHE), Health Protection Scotland, Public Health Wales, 2023). However, in developing countries, CDI epidemiology is underestimated due to limitations in surveillance protocols and diagnostic resources (Curcio et al., 2019b).

There are over 800 recognized strain types (ribotypes) of *C. difficile* but only toxin-producing strains are associated with disease (Dayananda and Wilcox, 2019). A wide array of virulence factors, such as surface proteins and toxins, influence the clinical manifestations of CDI. Toxins disrupt the epithelial gut barrier and induce inflammation, increasing submucosal edema, tissue damage and immune cells infiltration (Czepiel et al., 2019; Hernández Del Pino et al., 2021). All virulent strains carry a 19.6 kb pathogenesis locus (PaLoc) encoding *tcdR, tcdB, tcdE, tcdA,* and *tcdC* genes. PaLoc regulates the production of enterotoxin A (TcdA) and cytotoxin B (TcdB). Some ribotypes also encode the *C. difficile* transferase (CDT) binary toxin, another important virulence factor.

In the last two decades, the emergence of *C. difficile* strains has been associated with outbreaks in the epidemiology of CDI, leading to increased recurrence, time of hospitalization and mortality rates. First recognized in 2002, *C. difficile* BI/NAP1/027 clones have caused major epidemics throughout the developed world with substantial morbidity and mortality (Martin et al., 2016). In Argentina, the presence of this strain was reported by Cejas and collaborators in 2018 (Cejas et al., 2018).

Multiple strains can be detected by methods that can distinguish individual strains. In the context of *C. difficile*, these methods include restriction enzyme analysis (REA), pulsed field gel electrophoresis (PGFE), PCR ribotyping, multilocus variable number tandem repeat analysis (MLVA), multilocus sequence typing (MLST), and whole genome sequencing (WGS) (Dayananda and Wilcox, 2019). With the advent of next-generation sequencing, WGS is increasingly used as a fingerprinting method, allowing for accurate transmission tracking and providing a tool to identify and help control outbreaks (Cho et al., 2021). WGS can also detect genes that might contribute to identify virulence factors and new targets for therapies.

In this study, we performed WGS of a NAP1 commercial strain. We additionally designed functional assays to explore the ability of this strain to form resistance structures. We predicted vancomycin and tetracycline (commonly used antibiotics for CDI treatment) resistance genes and we also identified key factors for bacterial pathogenesis such as the presence of a type IV secretory system (T4SS) gene. These results allow a deep characterization of this NAP1 strain which is key for studies of host pathogen interaction.

## Materials and Methods

### Bacterial growth and DNA extraction

*Clostridioides difficile* derived from CDC20121308. Isolation: NY 2011. KWIK-STICK N° 01165P from Microbiologics^®^. CDC20121308 *C. difficile* strain was grown in CHROMagar^TM^ *C. difficile* (CHROMagar^TM^) in anaerobic conditions using Anaeropack® (Mitsubishi Gas Chemical America) at 37°C. One isolated colony was picked and suspended in brain heart infusion (BHI) (Oxoid) broth culture supplemented with 5% yeast extract and 0.1% cysteine (BHIS). After incubation in agitation for 48h under anaerobic conditions, 1 ml of culture was used for DNA extraction with EasyPure® Bacteria Genomic DNA Kit following the manufacturer’s instructions (Transgen Biotech). DNA integrity and concentration were measured by 1% agarose gel electrophoresis and Qubit Fluorometric Quantification (Thermo Fisher Scientific).

### Genome sequencing and assembly

Whole genome sequencing was performed using the Illumina Novaseq 6000 platform (Novogene Corporation Inc.) with a paired-end 150 strategy.

The raw sequencing data was quality-trimmed with FastQC^1^ and Trimmomatic (Bolger et al., 2014). The novo genome assembly was performed by SPAdes (Bankevich et al., 2012). Annotation was performed using Prokka (Seemann, 2014) and Rapid Annotation Subsystem Technology (RAST) (Aziz et al., 2008). 105 contigs were found.

The complete genome sequence of CDC20121308 *C. difficile* strain has been deposited on the National Center for Biotechnology Information (NCBI) database. The BioProject, BioSample, and RefSeq assembly accession numbers are PRJNA1054480, SAMN38922876 and GCF_035011885.1, respectively.

Annotation was performed using PGAP (Li et al., 2021), Prokka (Seemann, 2014) and Rapid Annotation Subsystem Technology (RAST) (Aziz et al., 2008).

### Whole genome analysis

Multi locus sequence type (MLST) was determined as previously reported (Griffiths et al., 2010) using public databases for molecular typing and microbial genome diversity (PubMLST)^2^. For comparative genome analysis, NAP1/BI/027 *C. difficile* strains R20291, 2007885, CD196 and BI1 (NCBI accession numbers: NZ_CP029423, NC_017178.1, NC_013315.1 and NC_017179.1, respectively) available in NCBI database were selected as references. Non-redundant nucleotide collection database from NCBI were used to compare *TcdC* gene of CDC20121308 against 630 strain.

Multiple Genome Alignment for genome comparison between the different strains was performed with MAUVE software (Darling et al., 2004) using progressive MAUVE alignment option. With R20291 strain as reference, draft genome contigs of CDC20121308 were ordered using MAUVE contig mover. Taking into account 70 and 60 percent as coverage and identity parameters, orthologues genes were obtained from MAUVE orthologues and used to generate Visualizing Intersecting Sets (UpSet^3^) to determine unique and shared genes between strains. For subsystems characterization, strains were annotated in Rapid Annotation using Subsystem Technology (RAST). R interface was used for representation of the data file (ggplot2 package) (Wickham, 2016).

Orthologous groups were identified using ProteinOrtho (Lechner et al., 2011) with the default settings, and used to generate Visualizing Intersecting Sets (UpSet^3^) to determine unique and shared proteins between strains. BLAST Ring Image Generator (BRIG) (Alikhan et al., 2011) and CIRCOS software (Krzywinski et al., 2009) were used for whole genome and specific locus comparison, respectively. Whole nucleotide sequence from R20291, CDC20121308, 2007855, CD196 and BI1 strains were used for align in BRIG. Selected genes enlisted in Supplementary Table 1 were used for the CIRCOS comparison.

### Colonial Morphology

Spread plate technique was used to grow CDC20121308 *C. difficile* strain in CHROMagar *C. difficile* (CHROMagar^TM^) for 48h in anaerobic conditions. The visual appearance and odor of the bacterial colonies were examined. Fluorescence under UV illumination was also corroborated. Gram staining using crystal violet and iodine was performed after fixating bacteria in glass slides. The presence of endospores and morphologic characteristics were evaluated using an oil immersion objective of the DM750 optical microscope (Leica Microsystems).

### Motility assay

The motility of the CDC20121308 strain was tested as previously reported (Stabler et al., 2009; Groß et al., 2018) with minor modifications. CDC20121308 *C. difficile* strain was cultured in CHROMagar^TM^ *C. difficile* (CHROMagar^TM^) under anaerobic conditions at 37°C. A fresh single colony was stab-inoculated in the center of 0.175% semi-solid BHI-agar tubes and incubated in anaerobiosis at 37°C. Monitoring of the developed diffusion radius around the inoculation stab for the following hours was performed, taking pictures at 12, 24 and 48h post inoculation.

### Biofilm formation

CDC20121308 *C. difficile* strain was growth in CHROMagar *C. difficile* (CHROMagar^TM^) for 48h in anaerobic conditions. One isolated colony was picked and suspended in 5ml of BHIS broth. After ON culture, 1ml of 1:100 dilution was incubated in BHIS supplemented with 0.1M D-dextrose without agitation using 24 wells culture plates (JET BIOFIL). After 5 days, culture medium was carefully discarded and plates wells were washed twice with PBS 1X at room temperature. For biofilm staining, 0.2% filtered crystal violet was added for 30 minutes. Cristal violet was removed and the wells were washed twice with sterile PBS. Excess of dye was removed by washing with ethanol: acetone (1:1). Biofilm formation was evaluated by bright field microscopy using Zeiss Axiovert 40CFL inverted fluorescence phase contrast microscope with a Plan Apochromatic10x/0.25/∞ objective.

### Antibiotic resistance

The Comprehensive Antibiotic Resistance Database (CARD) (Alcock et al., 2020) and manual search were used to predict resistance genes in CDC20121308 strain. To compare the tetracycline gene insertion of CDC20121308 against R20291, genome sequence ARTEMIS tool was used (Carver et al., 2005).

Minimum inhibitory concentration for doxycycline (a tetracycline) and vancomycin was determined by broth microdilution assays in 96 wells plate. A bacterial suspension of *C. difficile* CDC20121308 strain, in the late exponential phase, was incubated with serial dilutions of antibiotics (64μg/ml to 0.5μg/ml and 32μg/ml to 0.25μg/ml for doxycycline and vancomycin, respectively) in anaerobic conditions for 48h. BHIS culture without antibiotics was used as control. *C. difficile* growth was determined by evaluating OD in a microplate reader at 600nm. Vancomycin resistance was defined according to the breakpoint reported by EUCAST in ≥ 2µg/ml (EUCAST, 2023). The doxycycline breakpoint is not established by EUCAST and/or CLSI (CLSI, 2023) and was defined in ≥ 8µg/ml according to published data (Schmidt et al., 2007).

## Results and discussion

### Sequencing, genome organization and comparative analysis

The phylogenetic diversity of *C. difficile* has allowed for the emergence of several epidemic strains in recent years (Buddle and Fagan, 2023). In particular, the ribotype (RT) 027 lineage was responsible for a 2001 North American epidemic, which spread to the UK, peaking in 2004–2006 (Rupnik et al., 2009; He et al., 2013; Buddle and Fagan, 2023). A deeper characterization of the different *C. difficile* RT could lead to a better understanding of *C. difficile* pathogenesis and host pathogen interaction. In this work we characterized the NAP1 commercial CDC20121308 *C. difficile* strain. To this end, we performed a whole-genome shotgun strategy. The draft genome was 4,116,167 bases in length and the G+C content of the genomic DNA was 28.56 mol%. RAST annotation showed a N50 of 128.749 and a L50 of 10. A total of 3,717 coding sequences (CDS) and 45 tRNAs were predicted by PGAP (Supplementary Table 2).

Pathogenesis is triggered by the production of one or a combination of TcdA, TcdB, and CDT present in the hypervirulent toxinotypes (Kuehne et al., 2010; Gerding et al., 2013; Aktories et al., 2017). TcdA and TcdB genes are located in the pathogenicity locus (PaLoc) together with three other elements *tcdR*, *tcdC* and *tcdE* that regulate the expression and secretion of bacteriotoxins. The presence of TcdB in CDC20121308 strain has been previously characterized by MALDI-TOF. We further analyzed the presence of RT 027 lineage markers in our sequencing data. BLASTn alignment against virulent 630 strain (RT 012) of *C. difficile* showed a single-nucleotide deletion at position 117 and an 18-bp deletion at position 330–347 in the tcdC gene of CDC20121308 strain (Supplementary Figure 1). These deletions result in an inactivating frame-shift mutation in the tcdC gene, a negative regulator of toxins production. Both mutations are characteristics for RT 027, but the point mutation results in a truncated version of the anti-sigma factor TcdC that leads to the hypervirulence of this RT with high toxin expression (Curry et al., 2007; Dupuy et al., 2008; Carter et al., 2011).

Open Reading Frames (ORFs) BlastP analysis showed that CDC20121308 harbors the four-gene insertion sequence for ThyA, DHFR, Sir2 family of protein deacetylases homologous and thiamine biosynthesis protein (ThiC) that disrupt the thymidylate synthetase (ThyX). ThyX is present in many other pathogenic bacteria and provides a potential drug target for antibiotic therapies due to its central role in thymidylate DNA biosynthesis and as an RNA-binding protein. The four ORFs are present in a characteristic insert of RT 027 (Knetsch et al., 2011; Robinson et al., 2014). Remarkably, the replacement of native *thyX* by *thyA* provides a selective advantage in archaea and bacteria, such as in RT 027 strains which contain ThyA enzymes conferring higher rates of genome replication *in vivo* (Escartin et al., 2008).

100% coverage and identity for the ORFs was found when BLASTp was performed for R20291 and CDC20121308 RT 027 strains of *C. difficile*. Indeed, we identified the same percentage for binary toxin gene (cdtA and cdtB) (Supplementary Table 3), further demonstrating that lineage markers of RT 027 are present in the strain under study.

By using Bidirectional Best Hits (BBH) strategy we performed a comparative analysis with other four full-length genomes (i.e. closed assemblies) available in GenBank/NCBI (2007855, BI1, CD196 and R20291) (Table 1), all RT 027, group BI, NAP1 and toxinotype III. With ProteinOrtho, we determined the presence of 3,510 orthologs among all the evaluated BI/NAP1/027 *C. difficile* strains (Figure 1A). We identified 19 orthologous genes among CDC20121308 and the R20291, 5 with 2007855 strain and none for BI1 and CD196 (Figure 1A). In addition, 60 genes were only expressed by the CDC20121308 BI/NAP1/027 *C. difficile* strain.

**Figure 1.**
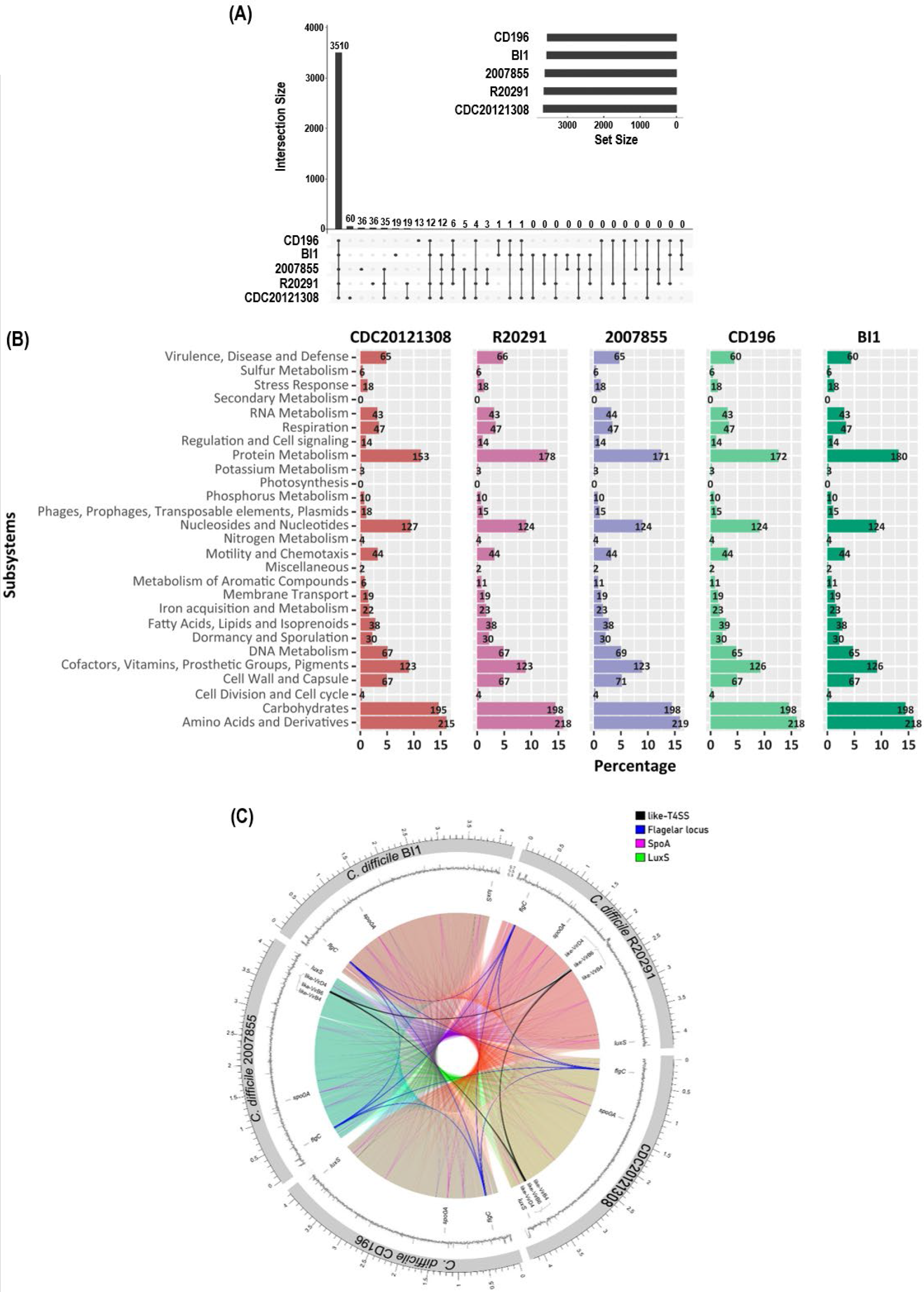
(A) Distribution of orthologous genes. Vertical bars show shared and unique coding sequences (CDS) between CDC20121308, CD196, BI1, 2007855 and R20291 strains. Horizontal bars indicate the CDS number of each strain used in this work. **(B) Comparison of RAST subsystem between strains.** CDS assigned to each subsystem category by RAST annotation database. **(C) CIRCOS software visualization comparison tool.** T4SS (black lines), flagellar (blue lines) and sporulation (magenta lines) loci as well as LuxS gene (green lines) were compared by CIRCOS. Background colors show homology between strains. Middle ring shows the GC content region.

**Table 1:** 027 Ribotypes strains used for genome comparison with CDC20121308. ST: sequence type. CDS: Coding sequence.

| Genome | Isolation<br>(year/country) | Accession number | RTo | Toxinotype | ST | Size (bp) | CDS | %GC |
| --- | --- | --- | --- | --- | --- | --- | --- | --- |
| CDC20121308 | 2011/NY US | GCF_035011885.1 | 027 | III | 1 | 4,116,167 | 3,717 | 28.6 |
| 2007855 | 2007/US | NC_017178.1 | 027 | III | 1 | 4,179,867 | 3,705 | 28.7 |
| R20291 | 2006/UK | NZ_CP029423.1 | 027 | III | 1 | 4,204,902 | 3,718 | 28.9 |
| BI1 | 1988/US | NC_017179.1 | 027 | III | 1 | 4,118,573 | 3,633 | 28.4 |
| CD196 | 1985/France | NC_013315.1 | 027 | III | 1 | 4,110,554 | 3,628 | 28.6 |

RAST annotation was performed to compare genes connected to subsystems and their distribution in different categories between the five strains (Figure 1B). RAST analysis showed a high similarity on the proportions of CDS assigned to each subsystem among all the strains. CDC20121308 showed the lowest proportion of CDS assigned to Protein metabolism subsystem. On the Virulence, disease and defense and the DNA metabolism subsystems, a higher proportion of CDS was observed in comparison to BI1 and CD196 strains (Figure 1B). Regarding PaLoc genes, we did not observe any differences between the five strains evaluated (data not shown).

The flag6ellar locus, Type IVC secretion system, master regulator of sporulation (SpoA) and luxS quorum sensing system of CDC20121308, 2007855, BI1, CD196 and R20291 *C. difficile* strains were compared using Circular Genome Data Visualization (CIRCOS) (Figure 1C). This analysis, together with RAST results, shows that the strain CDC20121308 evaluated in this work has higher similarity to the more recently isolated *C. difficile* strains R20291 and 2007855.

### CDC20121308 BI/NAP1/027 *C. difficile* strain characterization and functional assays

Of the total protein-coding genes found in our RAST annotation, 2,575 (68%) belongs to functional proteins, and the remaining genes were predicted as hypothetical proteins. Besides, CDC20121308 had 26% of their CDS assigned to subsystems (950 non-hypothetical and 32 hypothetical) and the remaining 74% (2,807 CDS, 1625 non-hypothetical and 1182 hypothetical) were not assigned to any subsystem. Regarding subsystems, a total of 273 were predicted in the *C. difficile* CDC20121308 genome using the RAST server, being the main subsystems amino acids and derivatives (16.04 %), carbohydrates (14.55 %), nucleosides and nucleotides (9.48 %), cofactors, vitamins, prosthetic groups, and pigments (9.18 %) (Figure 2).

**Figure 2.**
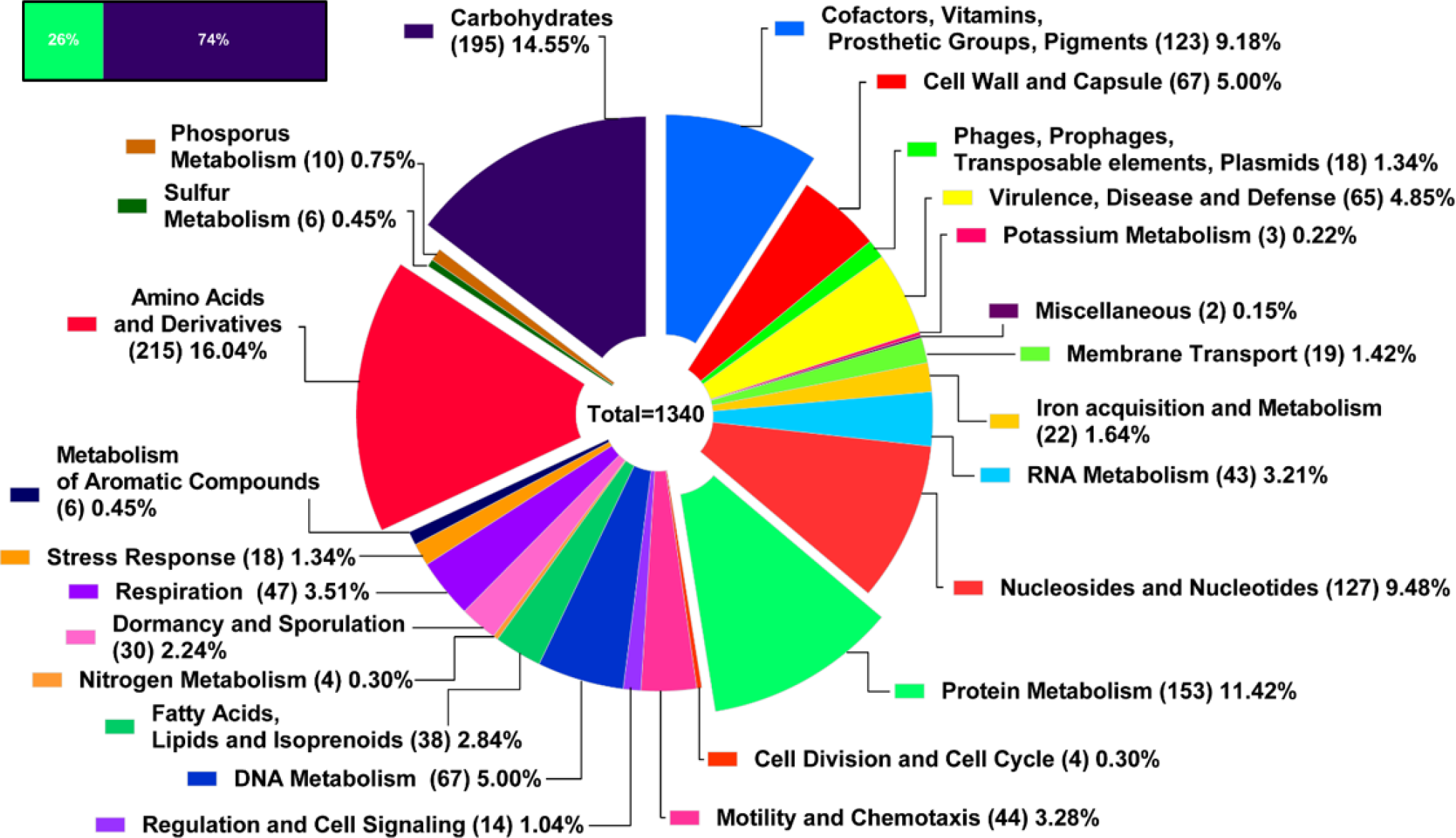
CDC20121308 subsystems. Proportion of CDS assigned (26%) or not assigned (74%) to subsystems by RAST tool. Pie chart shows the distribution of assigned CDS into each subsystem class.

To further characterize CDC20121308 BI/NAP1/027 *C. difficile* strain we performed functional assays. The colonies grown in CHROMagar *C. difficile* plates showed typical morphologic characteristics with road shape, irregular edges, slightly embossed and with brilliant surface (Figure 3A). Colonies of *C. difficile*, as usually described, also presented variations in size, they fluoresced under UV illumination and exhibited barn odor. Gram staining showed typical bacterial cell wall properties of *C. difficile* with gram-positive rods of different sizes and subterminal endospores (right panel).

**Figure 3.**
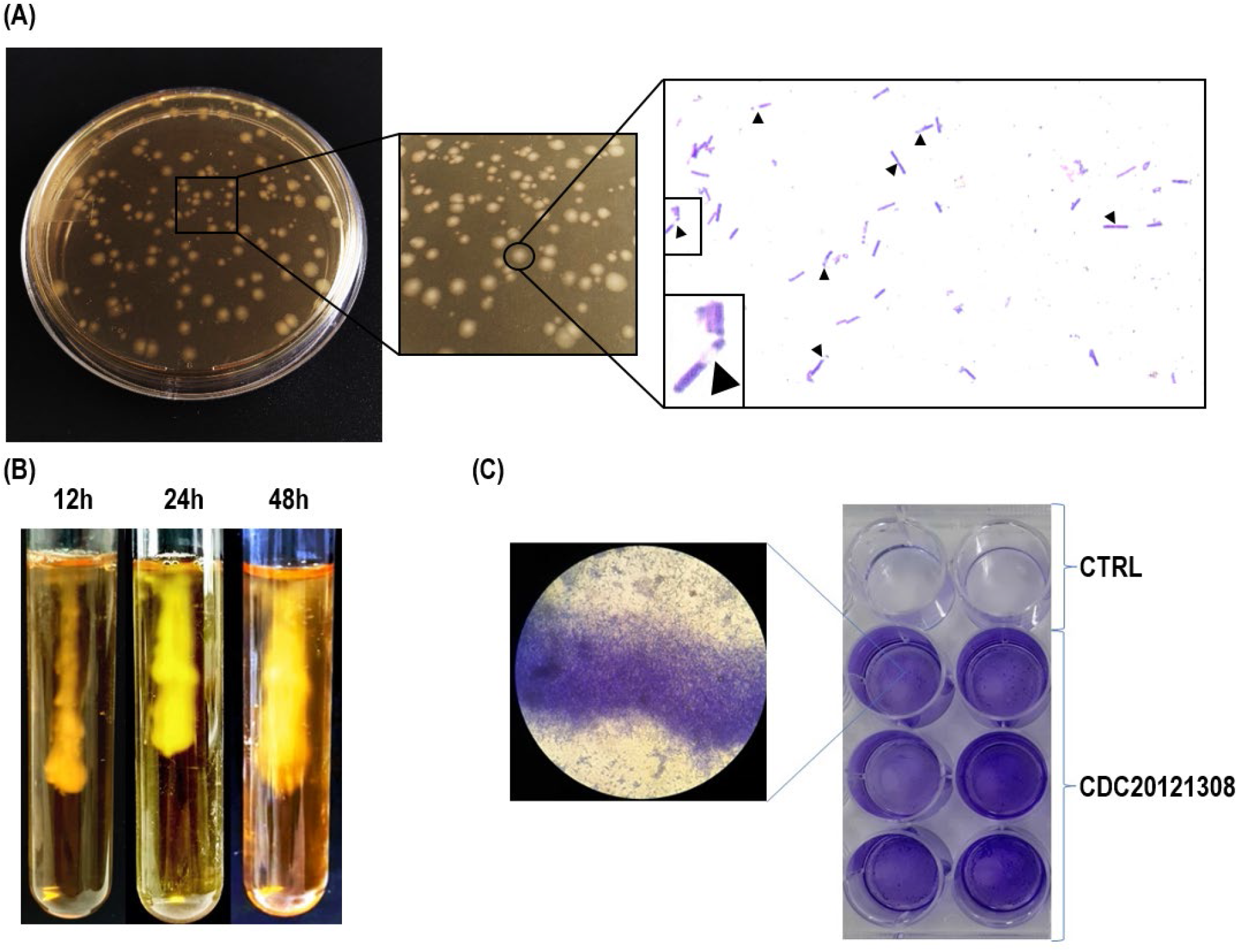
Functional and colony characterization of CDC20121308 strain. **(A)** CDC20121308 colonies were grown in CHROMagar^TM^ *C. difficile* medium for 48h. A magnification of the colonies is shown in the right upper image. Gram staining (right bottom image) was performed to check typical bacterial cell wall properties of *C. difficile*. Black arrowheads point endospores. **(B)** Motility assay was carried out in semisolid BHI agar and bacteria spreading was evaluated at 12, 24 and 48h. **(C)** Biofilm formation was assessed by crystal violet staining. CDC20121308 was incubated in BHIS with or without (Controls, CTRL) 0,1M D-dextrose. Microscopy image (left) shows biofilm adhered to the plate.

Some *C. difficile* surface proteins and regulatory factors key for biofilm formation such as flagella, pili, and Spo0A have been associated to gut surfaces adherence and pathogenicity during *C. difficile* infection (Abt et al., 2016; Frost et al., 2021). The majority of *C. difficile* strains, except 630, can change between an “on-off” state by sensing c-di-GMP levels, up-regulating or down-regulating flagella expression (Soutourina et al., 2013; Anjuwon-Foster and Tamayo, 2017; Arato et al., 2019). Our assays with the CDC20121308 BI/NAP1/027 *C. difficile* strain in soft agar tubes showed a diffuse growth away from the inoculum stab at 12, 24 and 48h, indicating a motile phenotype (Figure 3B). On the other hand, some reports consider flagella as an important mediator of biofilm production in late stages (Dapa et al., 2013) while others indicate that flagella expression is reduced when biofilm is formed (Maldarelli et al., 2016; Poquet et al., 2018). The 630 and the hypervirulent R20291 strains have the ability to form biofilm, although this capacity changes according to variations in the growth medium between both strains (Dapa et al., 2013). Moreover, *tcdB* expression levels (on R20291 strain) and sporulation (on 630 and R20291) has been positively correlated with the capacity of bacteria to produce biofilms (Maldarelli et al., 2016; Dawson et al., 2021). Our results showed that CDC20121308 BI/NAP1/027 *C. difficile* strain was also able to form biofilm (Figure 3C). Biofilm structures in the gut allow *C. difficile* persistence and could be related with the recurrent nature of CDI (Taggart et al., 2021). However, even though significant higher rates of sporulation were found in patients with recurrent CDI, no differences were observed for biofilm formation (Tijerina-Rodríguez et al., 2019). Our data, along with previous reports, suggest that biofilm formation is a complex and multifactorial strategy developed by *C. difficile* and seems to have no common pattern between strains. It is clear that biofilms confer greater survival to *C. difficile*, but whether they are necessary to increase strains virulence or whether they can be attributed to outbreaks or recurrences remains to be determined.

### Characterization of resistance genes to toxic compounds

By using CARD database, we predicted 11 genes for resistance to toxic compounds including the presence of efflux pump complex (cdeA) for disinfecting and DNA intercalating agents (Table 2). Moreover, the analysis showed the presence of several genes for resistance to Carbapenem, cephalosporines, macrolides (erythromycin, fidaxomicin), lincosamide (clindamycin), fluoroquinolones and glycopeptides (such as vancomycin) antibiotics (Table 2). Antibiotic resistance contributes to the emergence of hypervirulent strains, CDI outbreaks (He et al., 2013; Spigaglia, 2016; Peng et al., 2017) and reduces the efficiency of CDI treatment. The most common antibiotics used in CDI therapy are fidaxomicin, vancomycin and metronidazole (Nelson et al., 2017).

**Table 2:** CDC20121308 resistance genes against antibiotics, disinfecting agents and DNA intercalators.

| Resistance gene | Class | Drug class | Resistance mechanism |
| --- | --- | --- | --- |
| **b- lactamase** | **CDD-2** | **Carbapenem** | **Antibiotic inactivation** |
| **23s rRNA** | **Mutation** | **Macrolide antibiotic/**<br>**Lincosamide antibiotic** | **Antibiotic target alteration** |
| **Elongation factor Tu** | **Sequence variant** | **Elfamicyn antibiotic** | **Antibiotic target alteration** |
| **cdeA** | **Efflux pump complex** | **Disinfecting agents and intercalating dyes, acridine dye, fluoroquinolone antibiotic** | **Antibiotic efflux** |
| **gyrA** | **Amino acid substitution in gyrase subunit A** | **Fluoroquinolone antibiotic** | **Antibiotic target alteration** |
| **rpoC** | **Point mutation in the rpoC region** | **Glycopeptide antibiotic** | **Antibiotic target alteration** |
| **TetM** | **Tetracycline- resistant ribosomal protection protein** | **Tetracycline antibiotic** | **Antibiotic target alteration** |
| **sugE** | **Efflux transporter** | **Quaternary ammonium-resistance protein** | **Drug efflux pump** |
| **vanZ** | **Accessory protein** | **Teicoplanin resistance** | **Antibiotic target alteration** |
| **vanG** | **Accessory protein** | **Glycopeptide antibiotic** | **Antibiotic target alteration** |
| **vanR** | **Amino acid substitution (Thr115Ala)** | **Glycopeptide antibiotic** | **Antibiotic target alteration** |

Besides the commonly reported gene for vancomycin resistance, CDC20121308 contains a Thr115Ala substitution in the VanR gene (Figure 4A). We also found a resistance gene for tetracycline (*TetM*) which was absent in the other reference strains as shown by Mauve and Artemis alignment comparison (Figure 4B). TetM gene is the most common tetracycline resistance determinant that encodes a ribosomal protection protein which prevents the binding of tetracycline to 16S rRNA (Brown and Roberts, 1987; LaPlante et al., 2022). Many studies report that TetM gene is acquired within mobile genetics mobile elements or transposons by horizontal transfer (Dönhöfer et al., 2012; Dong et al., 2014; Imwattana et al., 2021; Lin et al., 2022) In CDC20121308, TetM was located in a 6,579 bp insertion together with others six genes, one of which corresponds to a macrolide-efflux protein transporter *(mefH* (Supplementary Table 4)). Tn5397, Tn916 and Tn6190 transposons often carry the *TetM* gene in *C. difficile* (Spigaglia et al., 2011; Imwattana et al., 2021). However, Tn6944 has recently been described to carry *TetM* and *mefH* genes together (Imwattana et al., 2021). According to our results, Tn6944 could be the transposon responsible for the CDC20121308 *C. difficile* strain insert.

**Figure 4.**
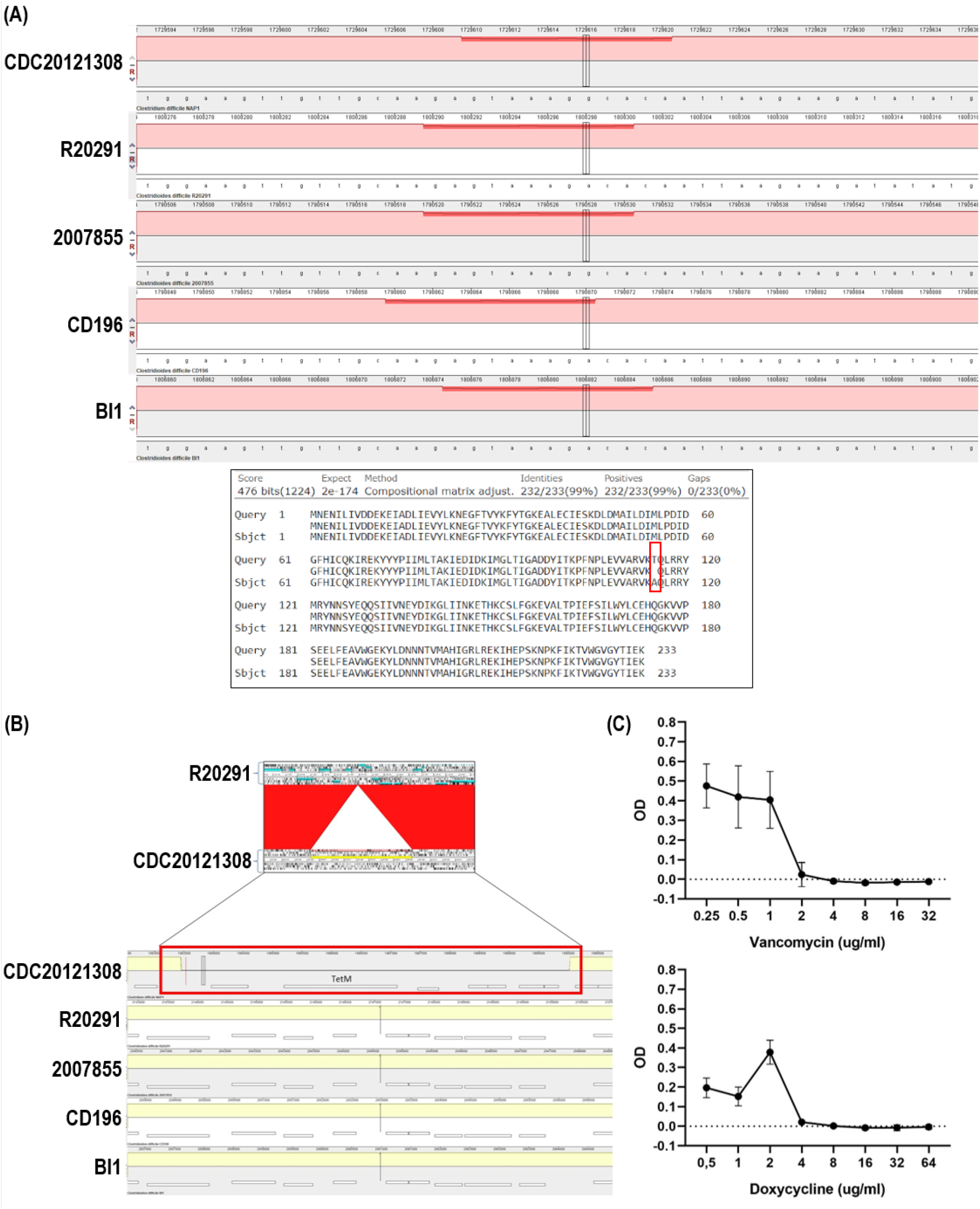
(A) *vanR* single mutation. A single mutation in *vanR* gene of CDC20121308 strain (upper image) was detected using MAUVE linear alignment. BLASTp of VanR protein sequence with a Thr115Ala substitution (red box) was identified using R20291 strain as reference (query). (**B) Tetracycline resistance gene**. Insertion carrying tetracycline resistance gene was observed by sequence alignment using Artemis comparison tool (upper image) against R20291 and MAUVE software (bottom image). **(C) Vancomycin and Doxycycline minimum inhibitory concentration (MIC).** Antibiotic resistance against vancomycin and doxycycline was determined by broth microdilution assay.

By broth microdilution assay we confirmed that the CDC20121308 BI/NAP1/027 *C. difficile* strain is resistant to Van and Tet with minimum inhibitory concentrations (MIC) of 4 μg/ml and 8 μg/ml, respectively (Figure 4C). Vancomycin is a first-line antibiotic used for CDI treatment (Peng et al., 2017). After completing initial antibiotic therapy, approximately 25% of patients develop CDI recurrence (Johnson et al., 2014; Gerding et al., 2018). Currently, a group of non-susceptible *C. difficile* strains have been identified (Peng et al., 2017; Darkoh et al., 2022). The VanA cluster, composed of *vanR*, *vanS*, *vanH*, *vanA*, *vanX*, *vanY*, and *vanZ*, mediates high-level vancomycin resistance (Darkoh et al., 2022). The full expression of the cluster leads to the remodeling of the peptidoglycan precursors (the target for vancomycin) decreasing the binding affinity of the drug more than 1000-fold (Miller et al., 2014; Darkoh et al., 2022). Moreover, *vanG* encodes for a d-alanine-d-serine ligase with a similar structure to that present in VanA gene cluster (Meziane-Cherif et al., 2012). In the CDC20121308 *C. difficile* strain, we found the *vanR*, *vanZ* and *vanG* genes. Our results indicate that CDC20121308 would belong to the vancomycin-resistant *C. difficile* strains group. These strains could lead to therapeutic failure and increased recurrence, highlighting the need for new therapeutic strategies.

Finally, using as reference the genome sequence of strain R20291, we generated a circular alignment on the BLAST ring imager (BRIG) (Figure 5). We identified areas with high and low identity among all the strains used in the study. The red rectangle indicates a zone of low identity with high GC content (Figure 5). With Mauve linear alignment (Figure 6) we identified that this region contains genes of the Type 4 Secretory System (T4SS). T4SS are a diverse superfamily of translocation systems to deliver DNA, protein and other macromolecules to bacterial or eukaryotic cell targets (Juhas et al., 2008; Sgro et al., 2019). T4SS plays a pivotal role in bacterial pathogenesis and the propagation of antibiotic resistance determinants throughout microbial populations (Ryan et al., 2023). Zhang and collaborators discovered a new class of T4SS, called Type IVC secretory system in gram-positive bacteria, including *C. difficile* (Zhang et al., 2012, 2017). Interestingly, they found that 8 of the 10 *C. difficile* strains studied have major components of the Type IVC secretion system. Of the four strains evaluated in our work only *C. difficile* CD196 and *C. difficile* BI1 did not carry a T4SS gene cluster (Zhang et al., 2017). Surprisingly, CDC20121308 has a cluster of genes on a genomic island (GI) of 58,079 bp that encodes some of the T4SS components (Supplementary Table 1). According to the classification of Zhang and collaborators, this GI corresponds to a new conjugative transposon 5-like (CTn5-like) (Zhang et al., 2017). Moreover, we identified VirD4, VirB4, and VirB6-like coding sequences similar to what was previously reported for the 630 strain (Sorokina et al., 2021). BLASTp of CDC20121308 VirD, VirB4 and VirB6-like protein sequences against *C. difficile* from NCBI database showed 100% homology with many strains, suggesting a conserved core of the type IVC secretory system (data not shown). R20291 shows more T4SS genes than CDC20121308 and 2007855 (Figure 5, Figure 6 and Supplementary Table 1).

**Figure 5.**
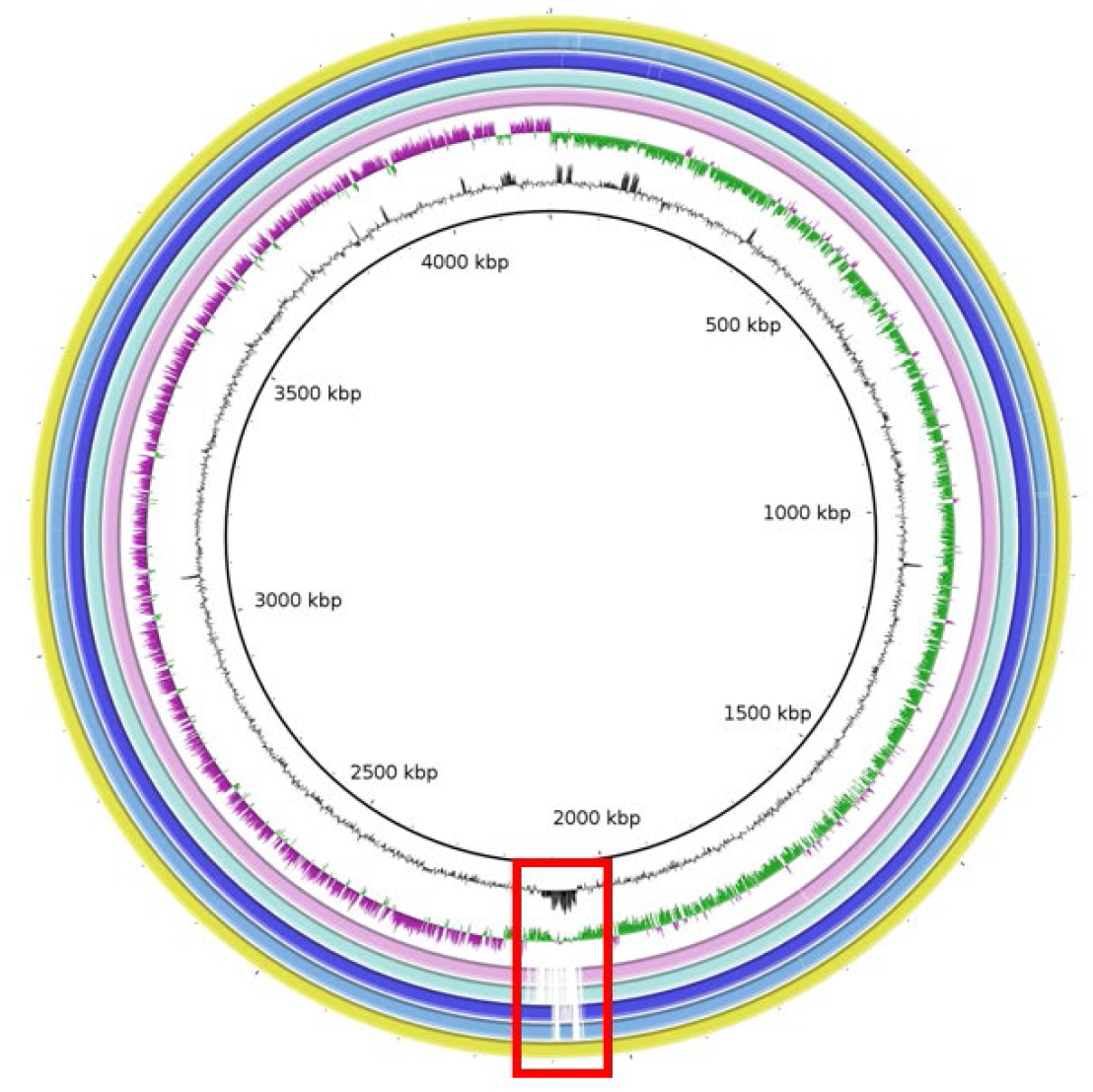
Circular alignment comparison. Rings represent strains genome alignment. From outer to inner, R20291, CDC20121308, 2007855, BI1, CD196, GC deviation (> 0%, purple; < 0%, green), GC content. White zone indicates low identity. R20291 strain was used as reference for BLAST. Red rectangle indicates low homology zone corresponding to T4SS genes.

**Figure 6.**
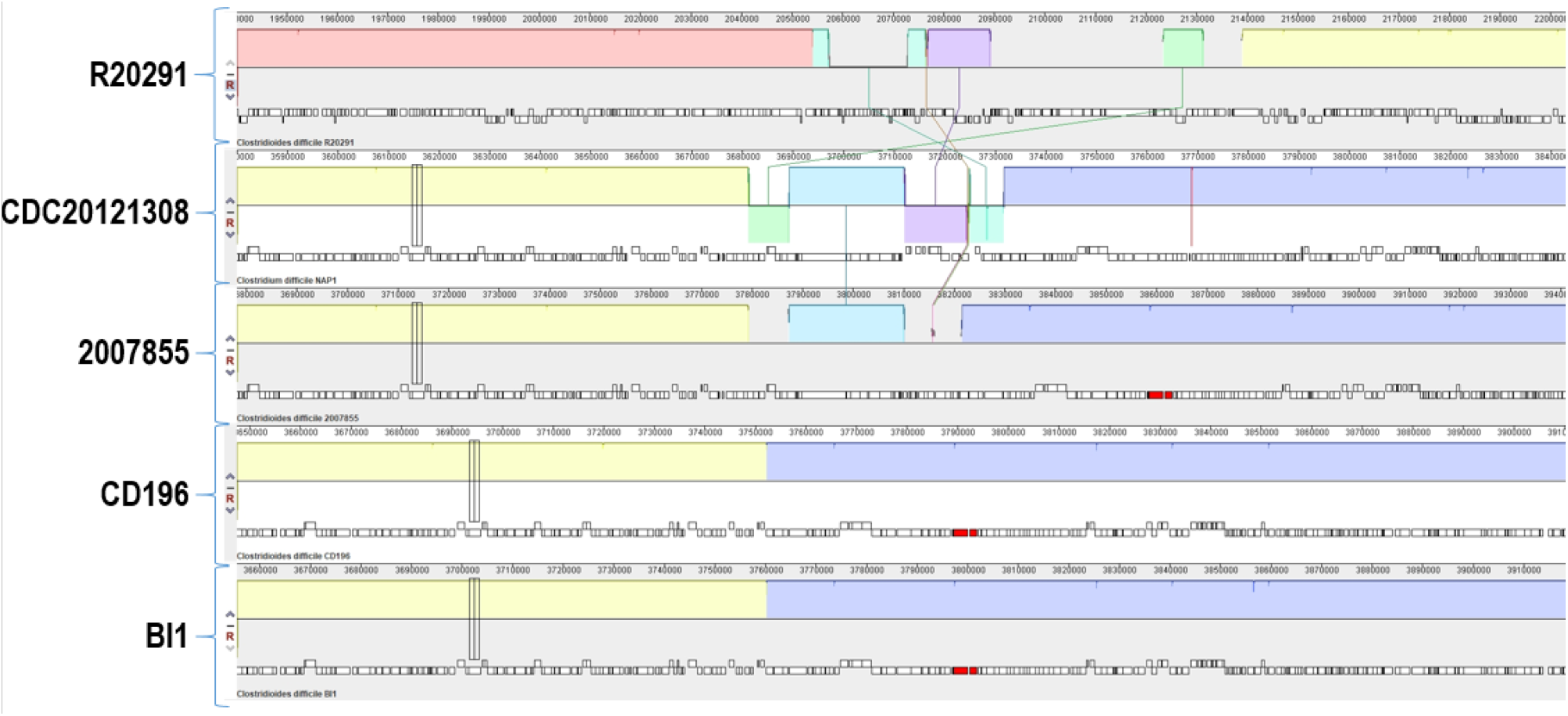
Linear alignment for T4SS components. T4SS insertion in R20291, CDC20121308 and 2007855 strains were detected by MAUVE alignment tool.

Altogether, these results show multiple characteristics related to resistance mechanisms that may contribute to the hypervirulence of the *C. difficile* CDC20121308 strain. The success of *C. difficile* as a pathogen is owed largely to its remarkable genome plasticity, allowing the acquisition of virulence factors and an array of resistance mechanisms (Buddle and Fagan, 2023). The interactions of *C. difficile,* an anaerobic pathogen, with the host and gut microbiota are highly intricate (Cheng and Unnikrishnan, 2023). Our study contributes to the evaluation of strains-dependent differences, which is necessary to establish good model systems for the study of CDI that facilitate the understanding of the disease *in vivo* and allow the search for new bacterial therapeutic targets.

## 2 Author Contributions

LAE, REHDP and VP conceived and designed the experiments. LAE and REHDP performed the experiments with AMB and SP contribution. LAE, MCP and FS analyzed the data. MCP, FS, DFDP and MAM helped with experimental design for sequencing, bioinformatics analysis and discussion of the results. DR contributed with the microbial assays and information about CDC20121308 strain. LAE, AMB, SP, REHDP and VP wrote the first draft of the manuscript. All authors contributed to manuscript reading and revising and approved the submitted version.

## 3 Funding

This work was supported by Universidad Nacional del Noroeste de la Provincia de Buenos Aires (grant numbers SIB 2113/2022 and “Proyectos de Investigación Interdisciplinarios de la UNNOBA” Res. CS 2190/2022, to V.P.). Agencia Nacional de Promoción Científica y Tecnológica, Fondo para la Investigación Científica y Tecnológica (ANPCyT-FONCyT, grant numbers PICT A PICT-2021-I-A-01119 to VP, PICT-2021-I-INVI-00584 to AMB, PICT-2021-I-INVI-00208 to SP and PICT-2021-GRFTI-00788). Consejo Nacional de Investigaciones Científicas y Técnicas (CONICET, grant number PIP 2021 11220200103137CO to VP and REHDP).

LAE was a doctoral fellow from Comisión de Investigaciones Científicas de la Provincia de Buenos Aires (CIC PBA), MCP and FS are doctoral fellows from CONICET, AMB and SP are postdoctoral fellows from CONICET. DR is staff member from ANLIS “Dr. Carlos G. Malbrán”. MAM, DFDP, REHDP and VP are researchers from CONICET.

## 4 Acknowledgments

We thank Natalia Menite, Gastón Villafañe and Lucia Romano for their technical assistance. We acknowledge Monica Machaín for their help with *C. difficile* microbiological culture.

## 5 Footnotes

1. https://bioinformatics.babraham.ac.uk/projects/fastqc

2. https://pubmlst.org/organisms/clostridioides-difficile

3. https://upset.app/

