## Supplementary Material for "Genome sequence and characterization of a hypervirulent BI/NAP1/027 *Clostridioides difficile* (CDC20121308)"

* These authors share first authorship

*** Correspondence:**

Pasquinelli, Virginia

**Supplementary Table 1. Selected genes for the CIRCOS comparison.**

*Attached excel as “Supplementary Table 1”*

**Supplementary Table 2. Genome overview of CDC20121308 based on Prokka and RAST annotations.**

| **Features** | **CDC20121308**  **PGAP** | **CDC20121308**  **Prokka** | **CDC20121308**  **RAST** |
| --- | --- | --- | --- |
| **Genome size** | **4,116,167** | **4,116,167** | **4,116,167** |
| **GC content (%)** | **28.56** | **28.56** | **28.56** |
| **Contigs number** | **105** | **105** | **105** |
| **CDSs** | **3,717** | **3,657** | **3,789** |
| **Total RNAs** | **61** | **56** | **56** |
| **tRNAs** | **45** | **48** | **47** |
| **rRNAs** | **16** | **8** | **9** |
| **Hypothetical proteins** | **379** | **1,558** | **974** |

*CDS: Coding sequence.

**Supplementary Table 3. BLASTp of RT 027 lineage markers.**

|  |  | **CDC20121308** | |
| --- | --- | --- | --- |
| **RT 027 lineage markers** | **R20291 protein ID** | **Identity (%)** | **Coverage (%)** |
| **ThyA (276aa)** | **WP_009887841.1** | **100** | |
| **DHFR (146aa)** | **WP_009887849.1** | **100** | |
| **SIR2 family protein (458aa)** | **WP_009887851.1** | **100** | |
| **ThiC (366aa)** | **WP_009892546.1** | **100** | |
| **cdtA (463aa)** | **WP_009890822.1** | **100** | |
| **cdtB ( 876aa)** | **WP_009890823.1** | **100** | |

**Supplementary Table 4. Genes located in 6,579 bp insertion.**

*Attached excel as “Supplementary Table 4”*

**Supplementary Figure 1. TcdC BLASTn gene.** BLASTn of tcdC gene against 630 strain show 18bp and single deletion (red arrows) in CDC20121308 strain. Query: 630 strain. Subject: CDC20121308.
