## Supplementary figures and images for "Genome sequence and characterization of a hypervirulent BI/NAP1/027 *Clostridioides difficile* (CDC20121308)"

### Supplementary Figure 1

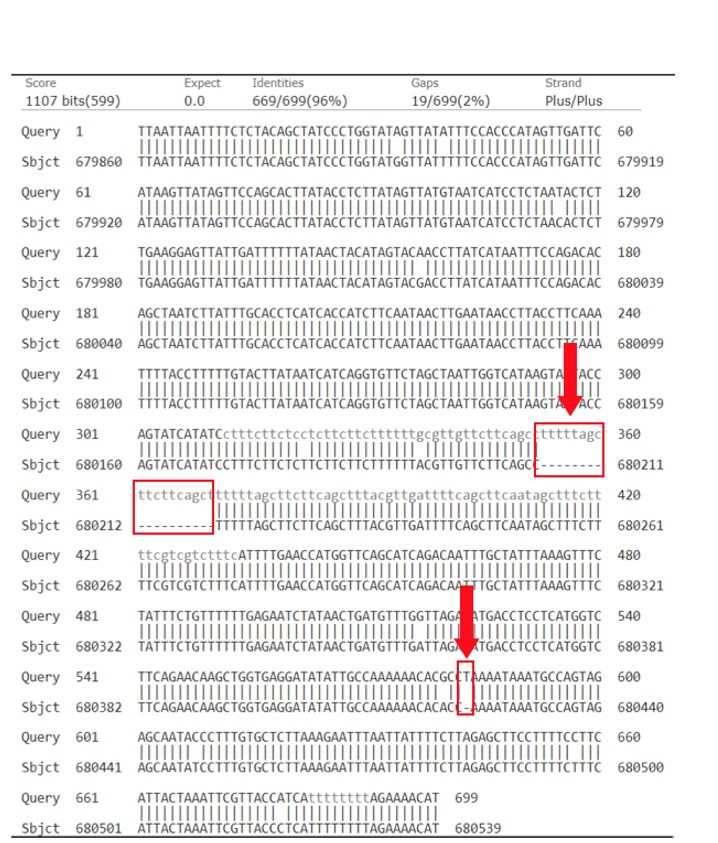
